# Aging imparts cell-autonomous dysfunction to regulatory T cells during recovery from influenza pneumonia

**DOI:** 10.1101/2020.06.05.135194

**Authors:** Luisa Morales-Nebreda, Kathryn A. Helmin, Nikolay S. Markov, Raul Piseaux, Manuel A. Torres Acosta, Hiam Abdala-Valencia, Yuliya Politanska, Benjamin D. Singer

## Abstract

Regulatory T (Treg) cells orchestrate resolution and repair of acute lung inflammation and injury following viral pneumonia. Compared with younger patients, older individuals experience impaired recovery and worse clinical outcomes after severe viral infections, including influenza and the novel severe acute respiratory syndrome coronavirus 2 (SARS-CoV-2). Whether age is a key determinant of Treg cell pro-repair function following lung injury remains unknown. Here, we show that aging results in a cell-autonomous impairment of reparative Treg cell function following experimental influenza pneumonia. Transcriptional and DNA methylation profiling of sorted Treg cells provide insight into the mechanisms underlying their age-related dysfunction, with Treg cells from aged mice demonstrating both loss of reparative programs and gain of maladaptive programs. Novel strategies that restore youthful Treg cell functional programs could be leveraged as therapies to improve outcomes among older individuals with severe viral pneumonia.

## Introduction

Age is the most important risk factor determining mortality and disease severity in patients infected with influenza virus or the novel severe acute respiratory syndrome coronavirus 2 (SARS-CoV-2) (*1, 2*). Global estimates of seasonal influenza-associated mortality range from 300,000-650,000 deaths per year, with the highest at-risk group comprised of individuals over age 75 (*3*). In the United States, influenza-associated morbidity and mortality have steadily increased, an observation linked to an expansion of the aging population. Pneumonia related to both severe influenza A virus and SARS-CoV-2 infection results in an initial acute exudative phase characterized by release of pro-inflammatory mediators that damage the alveolar epithelial and capillary barrier to cause refractory hypoxemia and the acute respiratory distress syndrome (ARDS) (*4*). If a patient survives this first stage, activation of resolution and repair programs during the ensuing recovery phase is crucial for restoration of lung architecture and function, which promotes liberation from mechanical ventilation, decreases intensive care unit length-of-stay and extends survival.

Immunomodulatory regulatory T (Treg) cells expressing the lineage-specifying transcription factor Foxp3 dampen inflammatory responses to both endogenous and exogenous antigens. Aside from their role in maintaining immune homeostasis through their capacity to suppress over-exuberant immune system activation, Treg cells reside in healthy tissues and accumulate in the lung in response to viral injury to promote tissue repair (*5, 6*). Our group and others have shown that in murine models of lung injury, Treg cells are master orchestrators of recovery (*7–10*). Treg cells are capable of promoting tissue regeneration and repair, at least in part through release of reparative mediators such as the epidermal growth factor receptor ligand amphiregulin (Areg), which induces cell proliferation and differentiation of the injured tissue (*11*).

Epigenetic phenomena, including DNA methylation, modify the architecture of the genome to control gene expression and regulate cellular identity and function throughout the lifespan (*12*). Aside from being one of the best predictive biomarkers of chronological aging and age-related disease onset, DNA methylation regulates Treg cell identity through tight epigenetic control of *Foxp3* and Foxp3-dependent programs (*13*). Biological aging is associated with a progressive loss of molecular and cellular homeostatic mechanisms that maintain normal organ function, rendering individuals susceptible to disease (*14, 15*). Because of their tissue-reparative functions, Treg cells are important modulators of the immune response that promotes tissue regeneration following injury (*16*). Whether age plays a key role in determining the pro-repair function of Treg cells in the injured lung during recovery from viral pneumonia remains unknown. If aging indeed impacts Treg cell-mediated recovery, is it a Treg cell-autonomous phenomenon or is it because the aging lung microenvironment is resistant to Treg cell-mediated repair? Using heterochronic (age-mismatched) adoptive Treg cell transfer experiments and molecular profiling in mice, we sought to determine whether the age-related impairment in repair following influenza-induced lung injury is intrinsic to Treg cells. Our data support a paradigm in which aged Treg cells fail to upregulate youthful reparative programs, activate maladaptive responses and consequently exhibit a cell-autonomous impairment in pro-recovery function, which delays resolution from viral-induced lung injury in aged hosts.

## Results

### Aging results in increased susceptibility to influenza-induced lung injury due to impaired recovery

To evaluate the age-related susceptibility to influenza-induced lung injury, we administered influenza A/WSN/33 (H1N1) virus via the intratracheal route to young (2 months) and aged (18 months) wild-type mice. Aged mice exhibited > 50% mortality when compared with young animals (**Figure 1A**), impaired recovery of total body weight following a similar nadir (**Figure 1B**) and more severe lung injury by histopathology at a late recovery time point, day 60 post-infection (**Figure 1C**). At this same time point, aged mice also displayed an increase in the total number of cells per lung (**Figure 1D**), which were mainly comprised of immune cells identified by the pan-hematopoietic marker CD45 (**Figure 1E**), suggesting non-resolving tissue inflammation during recovery in older mice. We next wanted to determine whether the age-related susceptibility to influenza-induced lung injury was due to a differential inflammatory response during the initial acute injury phase. Accordingly, we examined a different group of young and aged mice at a time point when viral clearance was complete (*17*) and weight nadir was observed in both groups, 14 days post-infection. Aged mice demonstrated increased mortality when compared with young animals at this time point (**Supplemental Figure 1A**), but other markers of acute inflammation, including weight loss (**Supplemental Figure 1B**), total lung cells (**Supplemental Figure 1C**) and total lung CD45^+^ cells in surviving animals were not significantly different between groups (**Supplemental Figure 1D**). Collectively, these results suggest that aging results in similar early injury but persistent lung inflammatory pathology during the recovery phase of influenza-induced lung injury.

**Figure 1.**
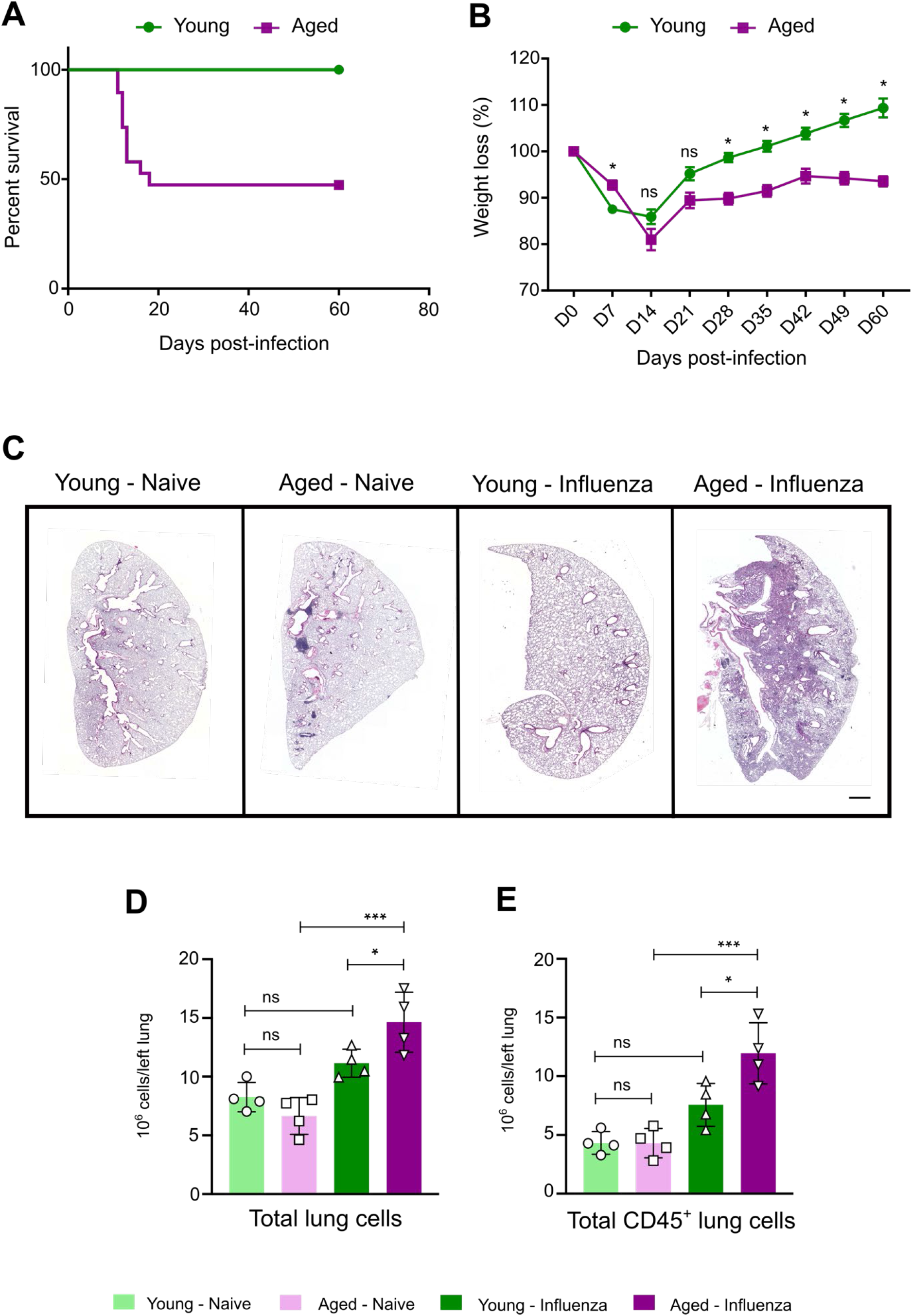
Aged mice demonstrate increased mortality and lung inflammation during recovery from influenza infection. (**A**) Survival curve of young (2 months, *n* = 20) and aged (18 months, *n* = 19) wild-type mice compared using the log-rank (Mantel-cox) test (*p* < 0.0002). (**B**) Weight loss percentage from baseline in young (2 months, *n* = 20) and aged (18 months, *n* = 19) mice compared using a mixed-effects model (REML) with Sidak’s *post-hoc* multiple comparisons test. Data presented as mean ± SEM. * *p* < 0.05, ns = not significant. (**C**) Representative lung histopathology (hematoxylin-eosin staining, scale bar = 1 mm) of young and aged mice during the naïve state and recovery phase following influenza infection (day 60). (**D**) Flow cytometry quantitative analysis of total number of cells and (**E**) Total number of CD45^+^ cells from left lung during the naïve state and recovery phase from influenza infection. Data presented as mean ± SD, *n* = 4 mice per group, one-way ANOVA with Holm-Sidak’s *post-hoc* testing for multiple comparisons. *** *p* < 0.0003, * *p* < 0.05, ns = not significant.

**Supplemental Figure 1.**
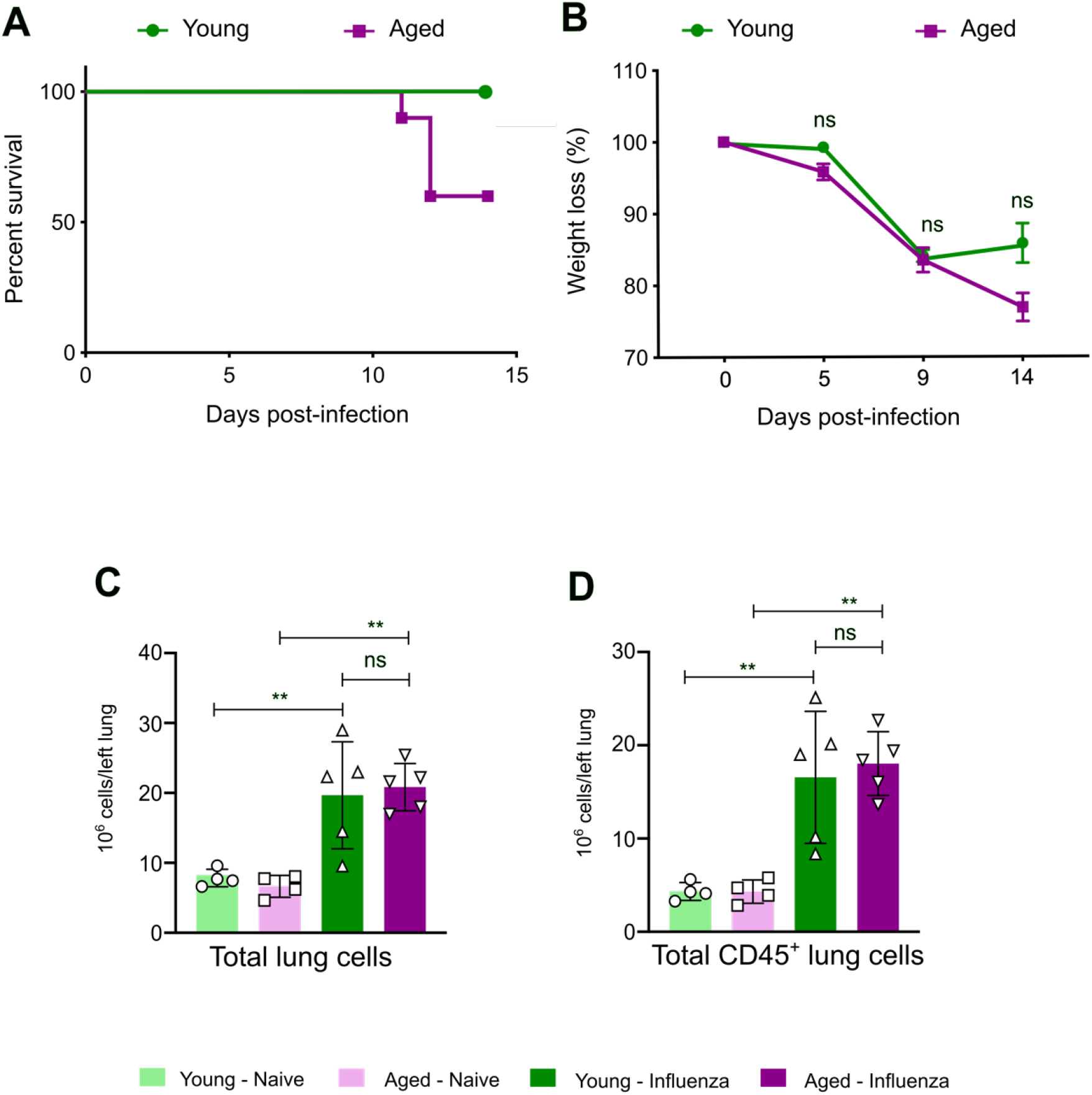
Aged mice demonstrate similar lung inflammation during early influenza-induced lung injury. (**A**) Survival curve of young (2-4 months, *n* = 10) and aged (18-22 months, *n* = 10) wild-type mice compared using the log-rank (Mantel-cox) test (*p* < 0.05). (**B**) Weight loss percentage from baseline in young (2-4 months, *n* = 10) and aged (18-22 months, *n* = 10) mice compared using a mixed-effects model (REML) with Sidak’s *post-hoc* multiple comparisons test. Data presented as mean ± SEM. ns = not significant. (**C**) Flow cytometry quantitative analysis of total number of cells and (**D**) Total number of CD45^+^ cells from left lung during the naïve state and early acute injury from influenza infection. Data presented as mean ± SD, *n* = 5 mice per group, one-way ANOVA with Holm-Sidak’s *post-hoc* testing for multiple comparisons. ** *p* < 0.005, ns = not significant.

### Aging results in deficient repair following influenza-induced lung injury

Having established that aging results in an increased susceptibility to persistent lung injury after influenza infection, we explored whether the impaired recovery in aged mice was linked to a persistent failure to repopulate the structural components of the alveolar-capillary barrier (i.e., failure to repair). Flow cytometry analysis (**Supplemental Figure 2**) of lung single-cell suspensions at day 60 post-influenza infection revealed an increased percentage of alveolar epithelial type 2 (AT2) cells (CD45^-^, T1α, CD31^-^, EpCAM^+^/CD326^+^, MHCII^+^) and endothelial cells (CD45^-^, T1α^-^, EpCAM^-^/CD326^-^, CD31^+^) (**Figure 2A-B** and **Figure 2D-E**, respectively), and a significant increase in the total number of AT2 and endothelial cells when compared with the naïve state (**Figure 2C** and **Figure 2F**, respectively).

**Figure 2.**
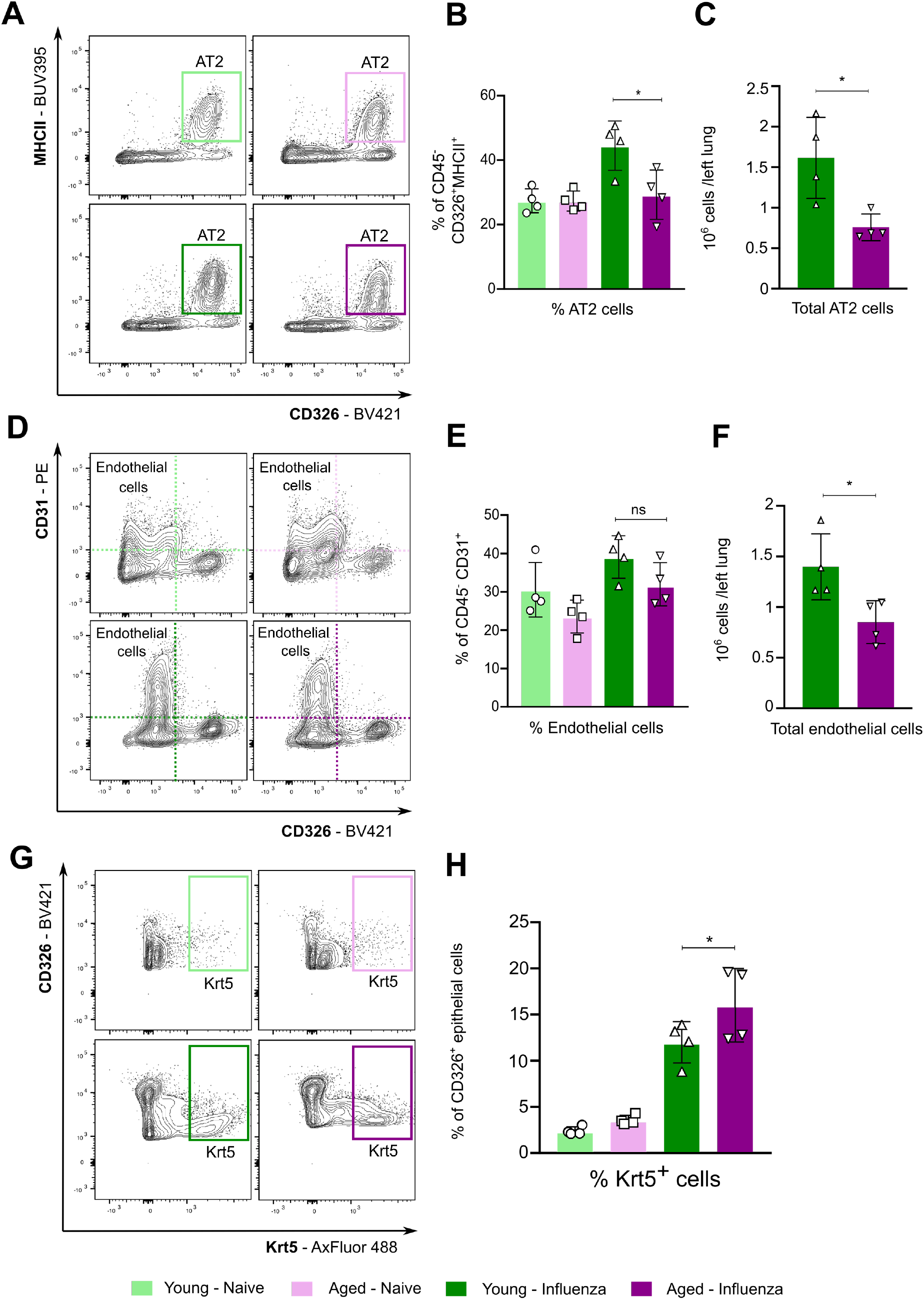
Aging results in failure to repopulate the alveolar epithelial-capillary barrier during recovery from influenza infection. (**A**) Representative flow cytometry contour plot analysis of type II alveolar epithelial cells (AT2). (**B**) Percentage of CD45^-^ CD326^+^ MCHII^+^ type II alveolar epithelial cells. (**C**) Total type II alveolar epithelial cells from left lung. (**D**) Representative flow cytometry contour plot analysis of endothelial cells. (**E**) Percentage of CD45^-^ CD31^+^ endothelial cells. (**F**) Total endothelial cells from left lung. (**G**) Representative flow cytometry contour plot analysis of Krt5^+^ cells. (**H**) Percentage of CD326^+^ Krt5^+^ epithelial cells. Data presented as mean ± SD, *n* = 4 mice per group, one-way ANOVA with Holm-Sidak’s *post-hoc* testing for multiple comparisons (B and E) or Mann Whitney test (C, F and H). * *p* < 0.05, ns = not significant.

In previous studies, investigators demonstrated that following influenza-induced lung injury, a population of cytokeratin 5^+^ (Krt5^+^) basal-like cells expand and migrate to the distal airspaces in an attempt to repair the injured epithelial barrier (*18*). These cells lack the capacity to transdifferentiate into functional AT2 cells, resulting in a dysplastic response that contributes to a dysregulated and incomplete repair phenotype following injury (*19*). Using a flow cytometry quantitative approach, we found that at 60 days post-infection, aged mice showed a significant increase in Krt5^+^ cells compared with young animals (**Figure 2G-H**). In summary, older mice failed to repair the injured lung during the recovery phase of influenza-induced lung injury.

**Supplemental Figure 2.**
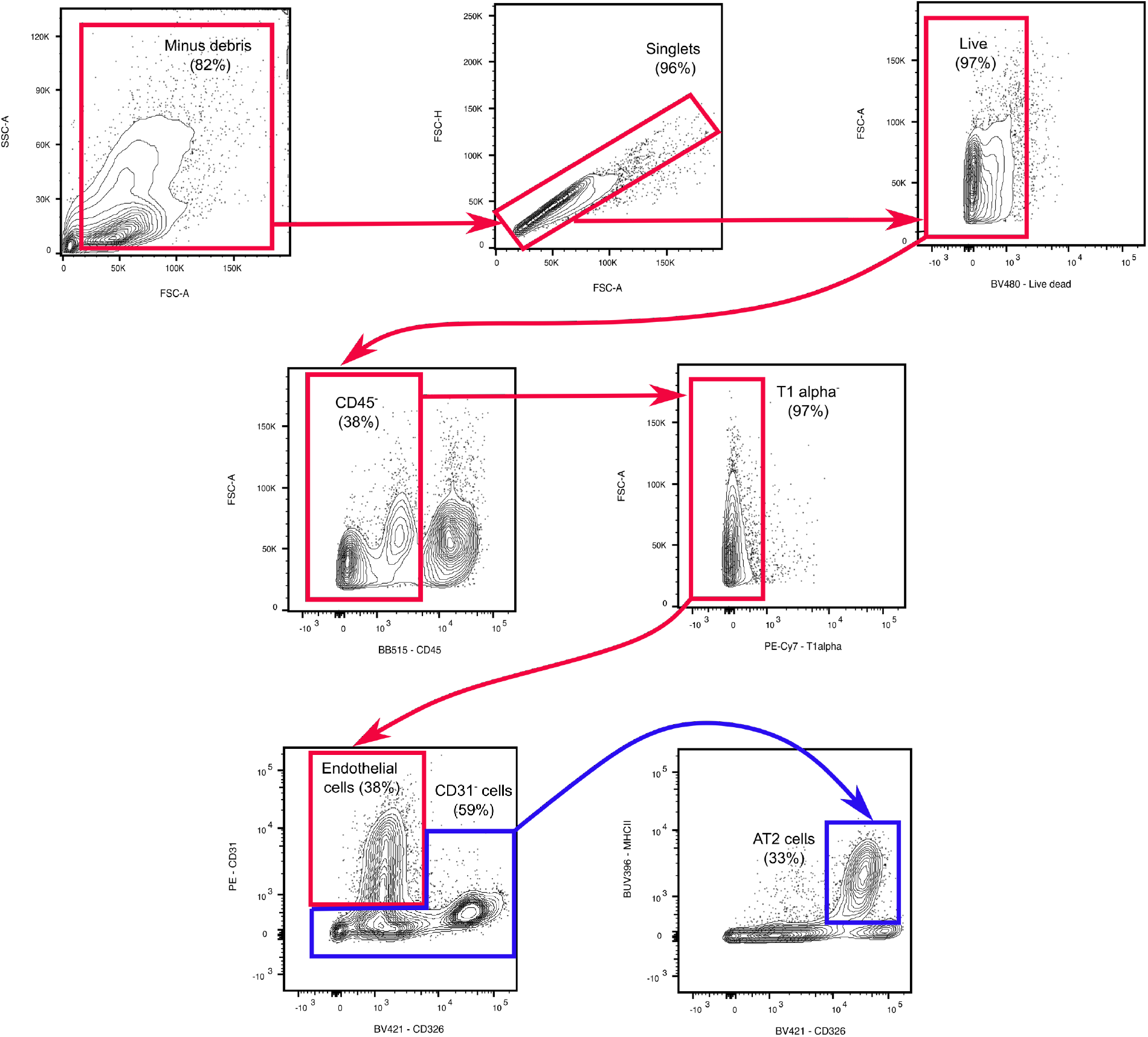
Sequential gating strategy used for flow cytometry analysis of type II alveolar epithelial cells and endothelial cells.

### Aging determines the pro-recovery function of Treg cells following influenza-induced lung injury

We and others have identified an essential role for regulatory T (Treg) cells in orchestrating resolution and repair of acute lung injury (*7–11*). Having established that aged mice fail to repair the injured lung, we next sought to determine whether this finding is due to age-related features altering the lung microenvironment or is driven by cell-autonomous, age-associated Treg cell factors. Thus, we performed heterochronic (age-mismatched) adoptive transfer of 1×10^6^ splenic young or aged Treg cells via retro-orbital injection into aged or young mice 24 hours post-infection (**Figure 3A**). Notably, adoptive transfer of young Treg cells into aged hosts resulted in improved survival when compared with aged mice that received PBS (control), while adoptive transfer of aged Treg cells into young hosts worsened their survival when compared with their respective controls (**Figure 3B**). We next turned to an inducible Treg cell depletion system using *Foxp3^DTR^* mice in order to eliminate Treg cells from recipients and specifically determine the age-related effect of donor Treg cells on the susceptibility to influenza-induced lung injury (**Figure 3C**). Adoptive transfer of aged Treg cells into Treg cell-depleted *Foxp3^DTR^* mice 5 days post-infection resulted in increased mortality when compared with adoptive transfer of young Treg cells (**Figure 3D**). Combined, our findings demonstrate that the loss of the Treg cell-associated pro-repair function in aged hosts is dominated by intrinsic, age-related changes in Treg cells and not conferred extrinsically by the aging lung microenvironment.

**Figure 3.**
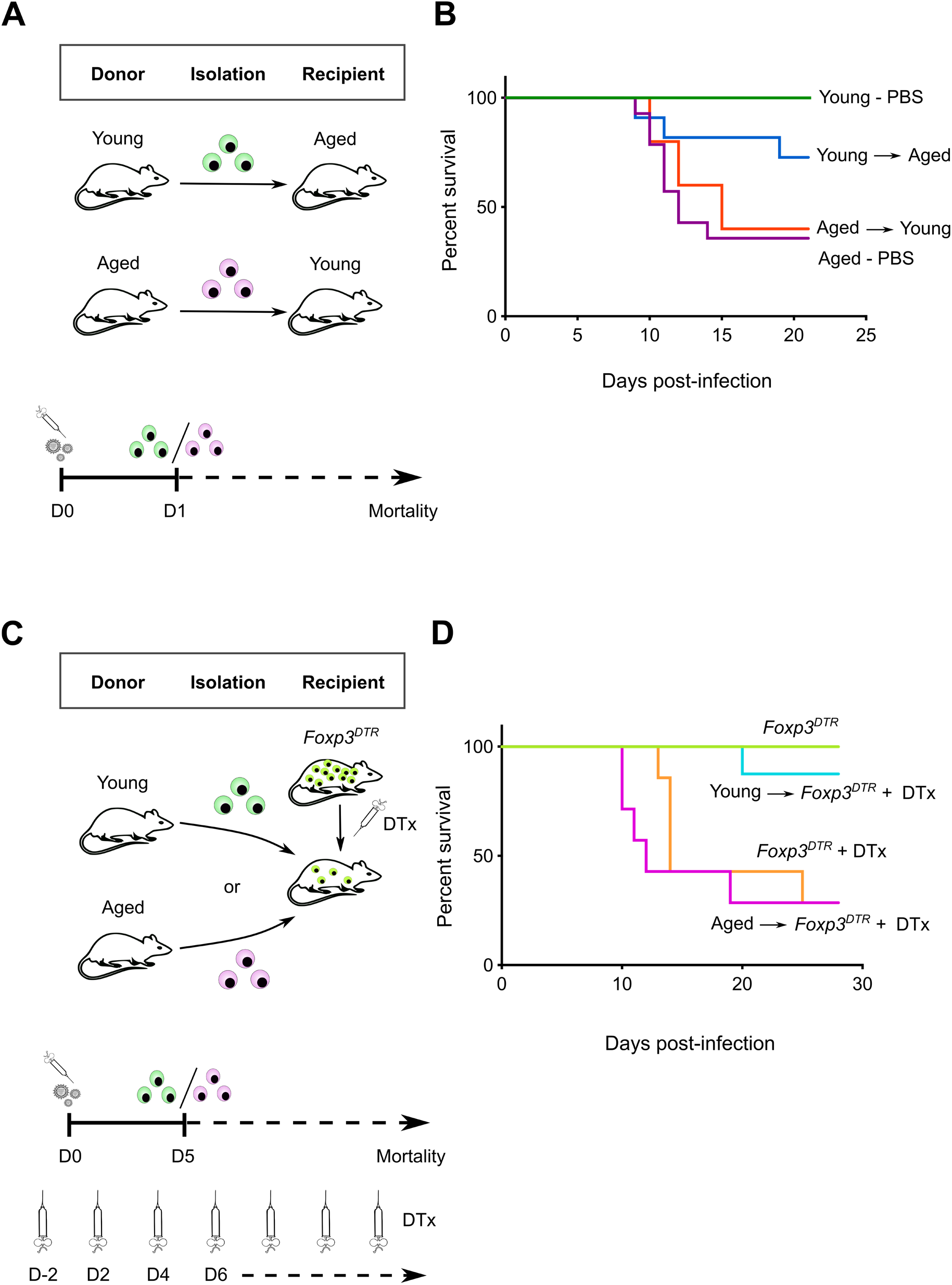
Age determines the tissue protective phenotype of Treg cells during influenza-induced lung injury. (**A**) Schematic of experimental design. (**B**) Survival curve of heterochronic adoptive Treg cell transfer experiments. *n* = 5-14 animals per group. (**C**) Schematic of experimental design. (**D**) Survival curve of heterochronic and isochronic adoptive Treg cell transfer experiments in *Foxp3^DTR^* mice. *n* = 7-8 animals per group except for the *Foxp3^DTR^* group (*n* = 3). DTx denotes diphtheria toxin.

### Aging results in the loss of pro-repair transcriptional programs in Treg cells during recovery from influenza-induced lung injury

To further explore the mechanisms underpinning the age-related loss of Treg cell pro-repair function following influenza infection, we performed gene expression profiling using RNA-seq on flow cytometry sorted lung Treg cells (**Supplemental Figure 3**) during the naïve state or late recovery phase from influenza (day 60 post-infection) (**Figure 4A**). Principal component analysis (PCA) of 3,132 differentially expressed genes (DEGs) after multiple group testing with false discovery rate (FDR) *q*-value < 0.05 demonstrated tight clustering by group assignment with PC1 reflecting the transcriptional response to influenza infection and PC2 reflecting age (**Figure 4B**). *K*-means clustering of these differentially expressed genes demonstrated that Cluster II was both the largest cluster and the one that defined the differential response to influenza infection between naïve and influenza-treated mice (**Figure 4C**). Notably, genes from this cluster were significantly upregulated among young Treg cells when compared with aged Treg cells following influenza infection (**Figure 4D**). Functional enrichment analysis revealed that this cluster was enriched for processes related to tissue and vasculature development and extracellular matrix formation (**Figure 4C**), suggesting a predominant reparative phenotype in young compared with aged Treg cells.

**Figure 4.**
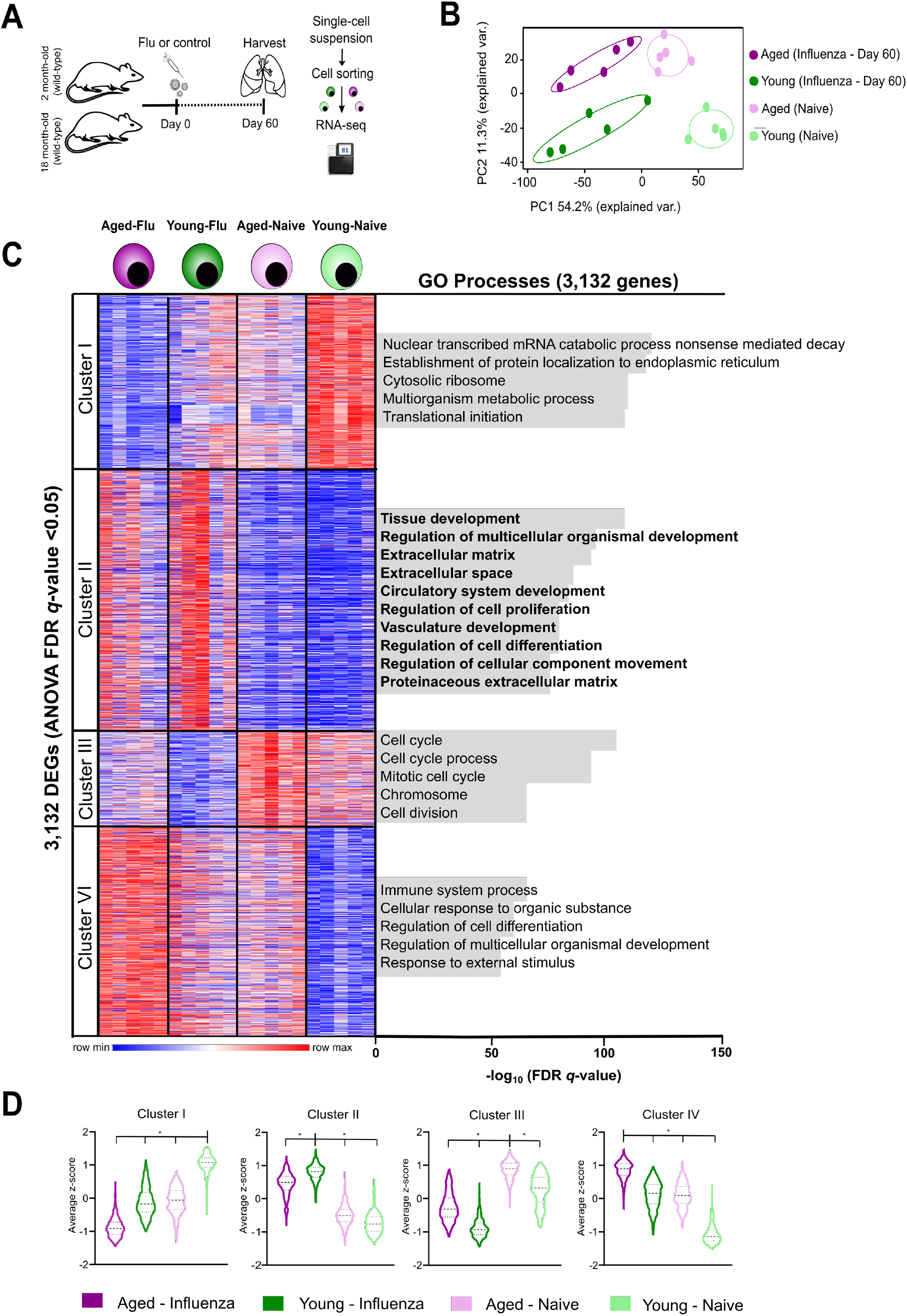
Young and aged Treg cells differ in their transcriptional response during recovery from influenza infection. (**A**) Schematic of experimental design. (**B**) Principal component analysis of 3,132 differentially expressed genes identified from a generalized linear model and ANOVA-like testing with FDR *q*-value < 0.05. Ellipses represent normal contour lines with one standard deviation probability. (**C**) *K*-means clustering of 3,132 genes with an FDR *q*-value < 0.05 comparing the cell populations from (B) with *k* = 4 and scaled as *z*-scores across rows. Top five gene ontology (GO) processes derived from clusters I, III and IV, and top 10 GO processes derived from cluster II are annotated and ranked by −log_10_-transformed FDR *q*-value. (**D**) Average *z*-scores for the four clusters shown in (C). Violin plots show median and quartiles. One-way ANOVA with two-stage linear step-up procedure of Benjamini, Krieger and Yekutieli with *Q* = 5%. * *q* < 0.0001.

**Supplemental Figure 3.**
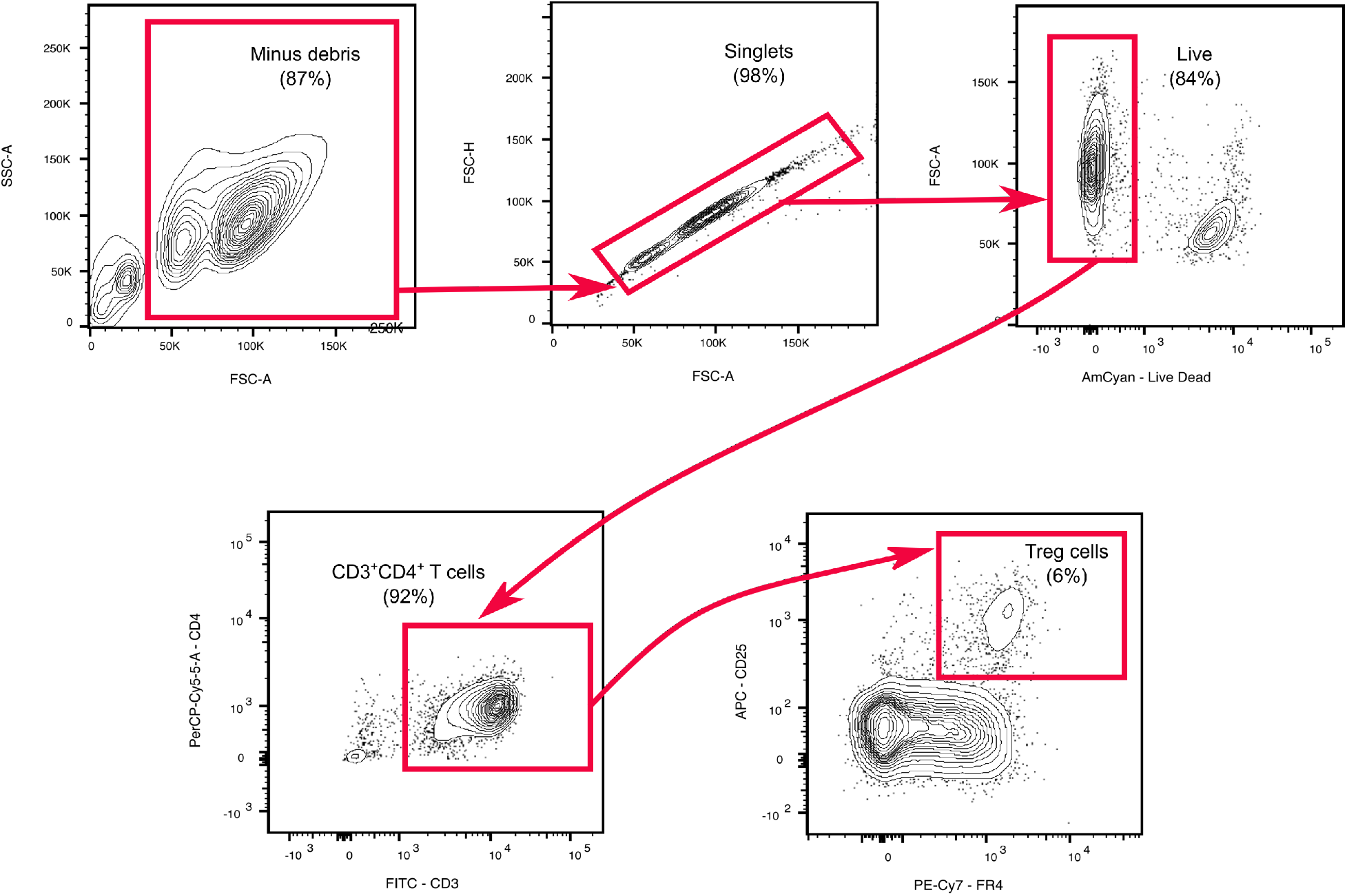
Sequential gating strategy used for flow cytometry sorting of Treg cells following enrichment for CD4^+^ T cells via magnetic bead separation.

We then performed an unsupervised analysis of the response to influenza infection from the naïve state to recovery phase between young and aged Treg cells. This analysis revealed upregulation of 1,174 genes (log_2_[fold-change] > 0.5, FDR *q*-value < 0.05) mostly linked to lung development (including epithelial and endothelial cell differentiation), extracellular matrix organization and wound healing (*Foxp2, Hhip, Klf2, Tns3, Hoxa5, Epcam, Erg, Bmper, Ereg, Lox, Tnc, Lama3* and *Spp1*) in young hosts (**Figure 5A-B**). We also found increased expression of genes associated with specialized Treg cell function in the maintenance of non-lymphoid tissue homeostasis and regenerative function (*Il1rl1* (encodes ST2), *Il18r1*, *Il10* and *Areg*). Aged Treg cells demonstrated increased expression of cell cycle genes (*Kif15* and *Cdk1*), neutrophil chemotaxis (*Cxcr1, Cxcl1* and *S100a9*) and cytotoxic effector function (*Gmzk*). Gene set enrichment analysis (GSEA) of the pairwise comparison between young and aged Treg cells during the recovery phase following influenza infection revealed that aged Treg cells downregulated repair-associated processes such as epithelial-to-mesenchymal transition, myogenesis and angiogenesis when compared with young hosts (**Figure 5C**). This pairwise comparison demonstrated that while young Treg cells exhibited significantly increased expression of genes associated with naïve (un-activated) state and lymphoid tissue markers akin to central Treg (cTreg) cell phenotype (*Lef1, Sell, Satb1, Bcl2, S1pr1, Gpr83* and *Igfbp4*), aged Treg cells upregulated genes implicated in effector Th1, Th17 and Tfr differentiation (*Tbx21/Cxcr3, Hif-1a* and *Sostdc1*, respectively), cell cycle (*Ccna2, Mmc3* and *Msi2*), T cell anergy (*Rnf128*) and DNA damage response (*Xrcc5* and *Rm1*) (**Figure 5C**). Collectively, these results reveal that while young Treg cells display a reparative phenotype during the recovery phase from influenza infection, aged Treg cells demonstrate both a less robust pro-repair phenotype when compared with young hosts and exhibit features of age-related maladaptive T cell responses, including effector T cell differentiation, cell cycle arrest and DNA damage responses (*20*).

**Figure 5.**
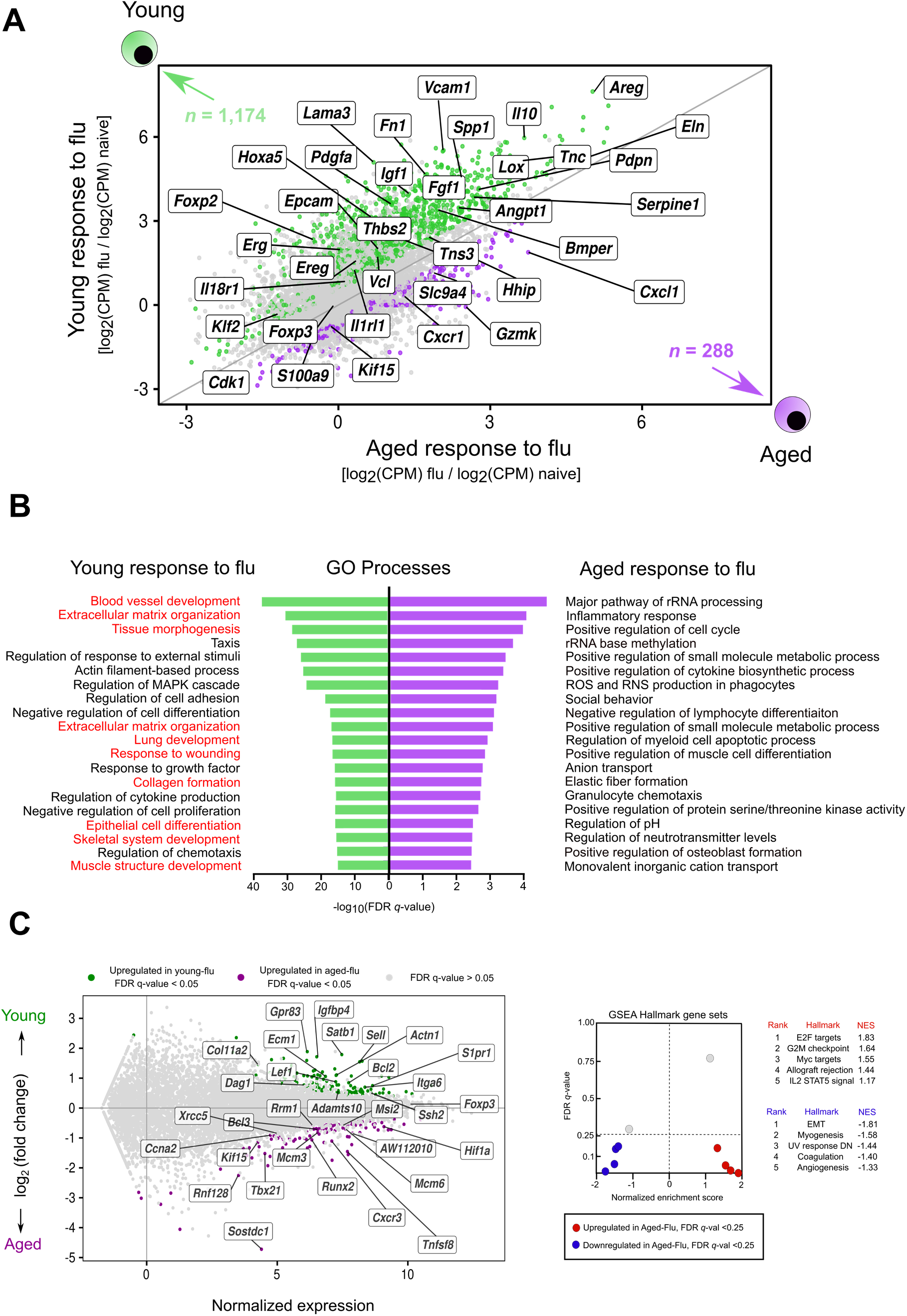
Young Treg cells upregulate a pro-repair transcriptional program during recovery from influenza infection. (**A**) Fold-change-fold-change plot for the young Treg cell response to influenza infection versus the aged Treg cell response to influenza infection highlighting genes exhibiting *q* < 0.05 and fold-change (FC) > 0.5 (green dots = young and purple dots = aged). Numbers of differentially expressed genes are indicated. (**B**) Top 20 gene ontology (GO) processes derived from differentially expressed genes (*q* < 0.05) from (A) for young Treg cells (1,174 genes) and aged Treg cells (288 genes) are annotated and ranked by −log_10_-transformed FDR *q*-value. Red font denotes pro-repair processes in young Treg cells. (**C**) MA plot comparing gene expression of young Treg versus aged Treg cells during the recovery phase from influenza infection. Genes of interest are annotated. (**D**) Gene set enrichment analysis (GSEA) plot highlighting key statistics (FDR *q*-value and normalized enrichment score or NES) and gene set enrichment per phenotype. Genes were ordered by log_2_(fold-change) and ranked by the aged Treg cell phenotype. Red dots denote gene sets with a positive enrichment score or enrichment at the top of the ranked list. Blue dots denote gene sets with a negative enrichment score or enrichment at the bottom of the ranked list.

### Regulatory T cells from aged mice demonstrate a Th1-like phenotype during recovery from influenza pneumonia

Upon stimulation, Treg cells can exhibit phenotypic and functional adaptability through upregulation of transcription factors and chemokine receptors akin to effector T helper cell subsets (e.g., Th1, Th2, Th17 and T follicular regulatory or Tfr) (*21*). The resulting effector Treg (eTreg) cells with Th-like phenotypes migrate to inflammatory non-lymphoid tissues and acquire transcriptional and functional programs that mirror the T effector responses they intend to suppress (*22, 23*). Having observed transcriptional upregulation of effector-associated factors in the aged Treg cell response during recovery from influenza infection, we next performed flow cytometry analysis to quantify some of the canonical effector-associated transcription factors and cytokines at the protein level. During the recovery phase following influenza infection, aged mice exhibited a higher percentage of eTregs when compared with young hosts (**Supplemental Figure 4A**). These aged eTreg cells demonstrated significant upregulation of the transcription factors Tbet and Ror-γt, which are canonical master regulators of Th1 and Th17 responses, respectively (**Supplemental Figure 4B**). Intracellular cytokine profiling of these cells also demonstrated a significant increase in production of Ifn-γ by aged Treg cells (**Supplemental Figure 4C**) when compared with young cells. Profiling of multiple well-established Treg cell suppressive markers, including *Foxp3*, showed no significant age-related difference during the recovery phase of influenza infection (**Supplemental Figure 5**). These data suggest that aged Treg cells mainly upregulate effector programs that, while protective during an insult such as influenza-induced acute lung injury, when observed during the recovery phase could represent an age-related maladaptive Treg cell response leading to unremitting lung inflammation and injury.

**Supplemental Figure 4.**
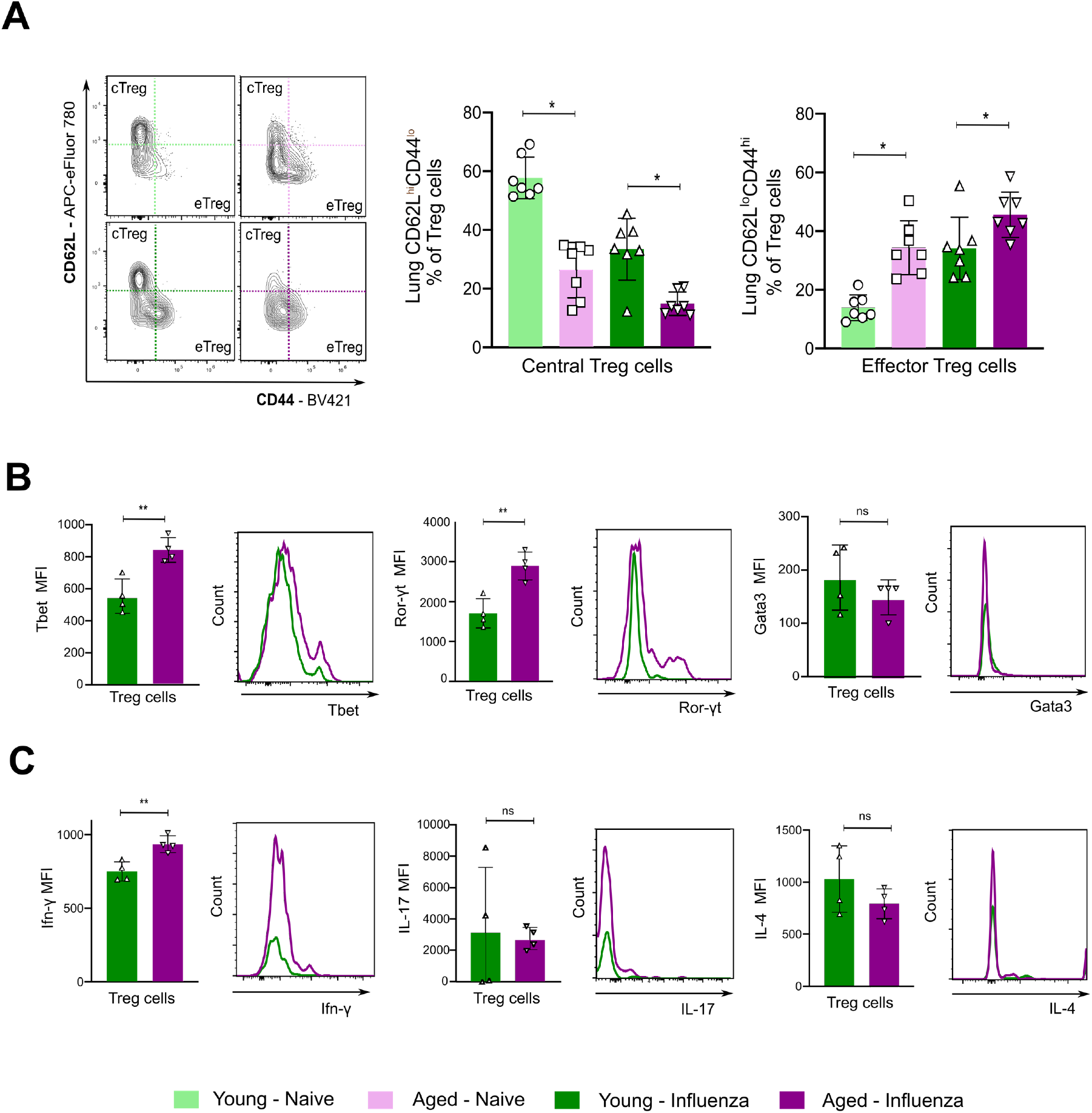
Aging results in induction of a Treg cell-Th1-like phenotype following recovery from influenza infection. (**A**) Representative flow cytometry contour plot analysis of central Treg (cTreg) cells and effector Treg (eTreg) cells. Percentage of cTreg cells (CD62L^hi^ CD44^lo^) and eTreg cells (CD62L^lo^ CD44^hi^) from sorted lung Treg cells (Supplemental Figure 3). (**B**) Representative histogram plots and pairwise comparison of median fluorescence intensity (MFI) from T helper cell canonical transcription factors. (**C**) Representative histogram plots and pairwise comparison of MFI from pro-inflammatory cytokine profiling. Data presented as mean ± SD, *n* = 4 mice per group, one-way ANOVA with Holm-Sidak’s *post-hoc* testing for multiple comparisons (A) or Mann Whitney test (B and C). ** *p* < 0.005, * *p* < 0.05, ns = not significant.

**Supplemental Figure 5.**
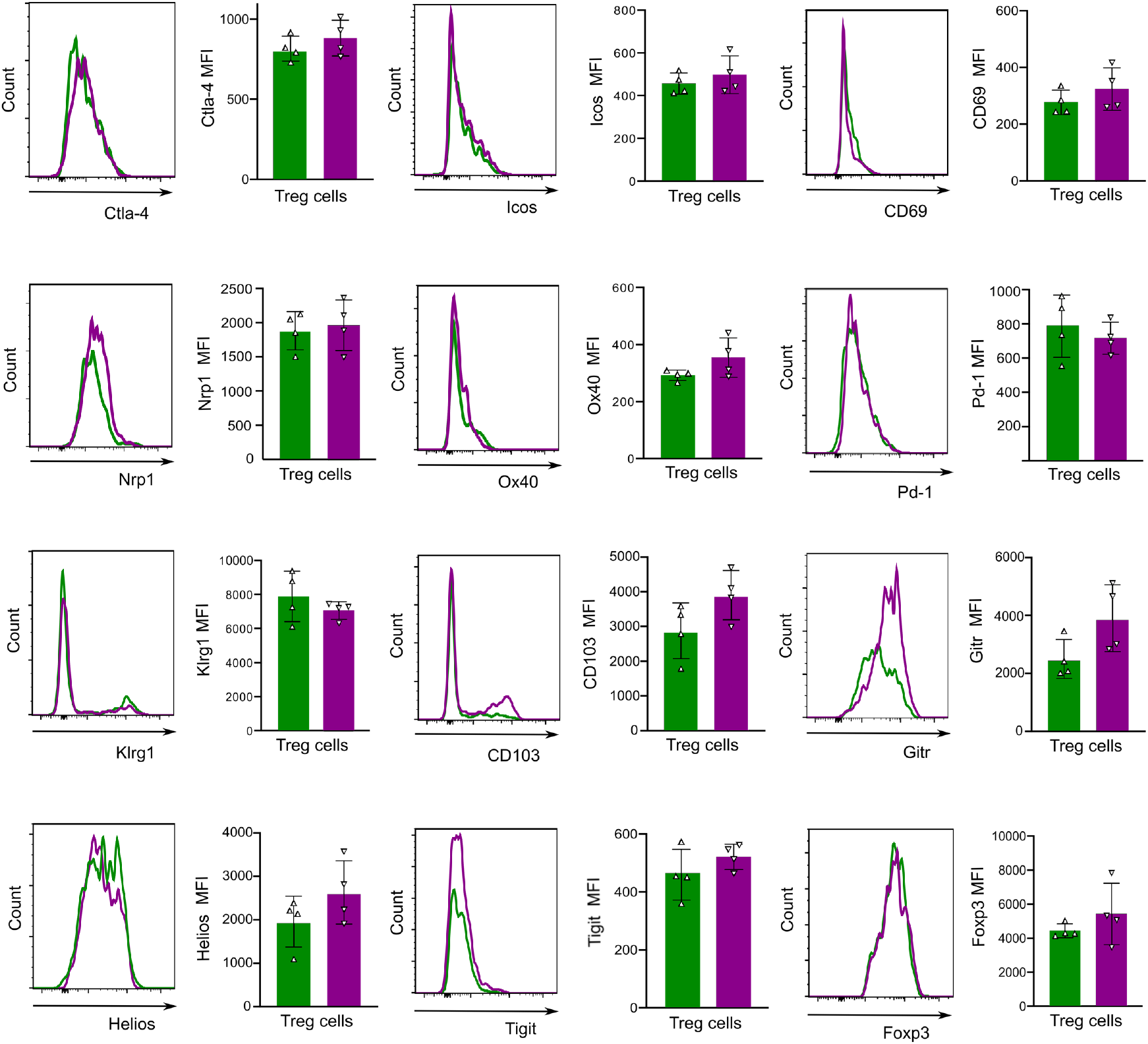
Aging does not result in changes in the suppressive phenotype of Treg cells during recovery from influenza infection. Representative flow cytometry histogram plots of Treg cell-associated activation and suppressive markers in lungs of young and aged mice harvested at day 60 post-influenza infection. Data presented as mean ± SD, *n* = 4 mice per group. All pairwise comparisons of each marker’s median fluorescence intensity (MFI) were not statistically significant by the Mann-Whitney test.

### Aged mice exhibit a less robust reparative response than young hosts during recovery from influenza pneumonia

To further investigate the age-related reparative function of Treg cells during the recovery phase of influenza infection, we performed pairwise comparisons of both young and aged Treg cells from the naïve state and recovery phase (day 60 post-infection) (**Supplemental Figure 6A-B**). We found that during the Treg cell response to influenza, there were 1,678 upregulated genes in young mice and only 445 upregulated genes in aged mice when compared with their respective naïve state (FDR *q*-value < 0.05). Gene set enrichment analysis revealed upregulation of pro-repair hallmark processes in both young and aged Treg cells during recovery from influenza infection (**Supplemental Figure 6C-D**). We next compared the age-related transcriptional response to influenza infection and found 342 shared genes between both age groups that were associated with pro-repair processes (**Supplemental Figure 6E**). The remaining 1,336 uniquely upregulated genes in young Treg cells were linked to reparative processes in contrast with the 103 uniquely upregulated genes in aged Treg cells. Altogether, these results suggest that while aged Treg cells show a reparative response during recovery from influenza infection, it is not as robust as the response exhibited by young Treg cells.

**Supplemental Figure 6.**
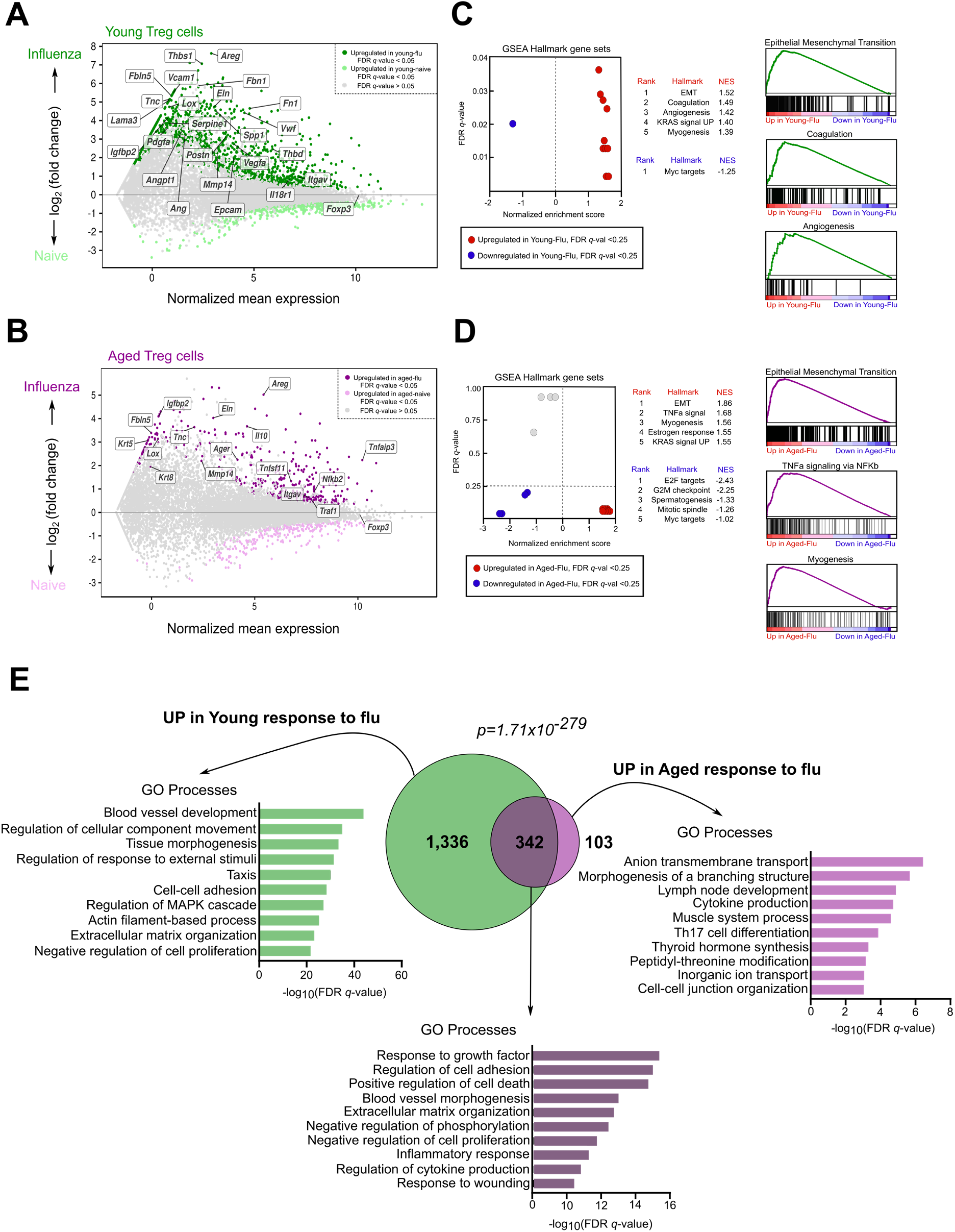
Aged Treg cells fail to upregulate a pro-repair transcriptional program to the same extent as young Treg cells during recovery from influenza infection. (**A**) MA plot comparing gene expression of young Treg cells during the naïve state from young Treg cells during the recovery phase from influenza infection. Genes of interest are annotated. (**B**) MA plot comparing gene expression of aged Treg cells during the naïve state from aged Treg cells during the recovery phase from influenza infection. Genes of interest are annotated. (**C**) and (**D**) GSEA plots highlighting key statistics (FDR *q*-value and normalized enrichment score or NES) and enriched gene sets per phenotype. Genes were ordered by log_2_(fold-change) and ranked by the young Treg cell-flu (**C**) aged Treg cell-flu (**D**) phenotype. Red dots denote gene sets with a positive enrichment score or enrichment at the top of the ranked list. Blue dots denote gene sets with a negative enrichment score or enrichment at the bottom of the ranked list. (**E**) Venn diagram partitioning into upregulated genes in young Treg cells during recovery from influenza infection (green circle, 1,336 genes), upregulated genes in aged Treg cells during recovery from influenza infection (light purple circle, 103 genes) and upregulated genes in both young and aged Treg cells during recovery from influenza infection (dark purple intersection, 342 genes). FDR *q*-value < 0.05. A hypergeometric *p*-value is shown. Top 10 gene ontology (GO) processes derived from genes in each partition of the Venn diagram are annotated and ranked by −log_10_-transformed FDR *q*-value.

### DNA methylation regulates the transcriptional pro-repair response to influenza infection

In addition to representing one of the hallmarks of aging, epigenetic phenomena such as DNA methylation regulate the development, differentiation and functional specialization of T cell lineages, including Treg cells (*12–14*). Therefore, we reasoned that age-related changes to the Treg cell DNA methylome could inform the divergent pro-repair transcriptional response seen between young and aged Treg cells following influenza infection. We performed genome-wide (5’-cytosine–phosphate–guanine-3’) CpG methylation profiling with modified reduced representation bisulfite sequencing (mRRBS) of sorted lung Treg cells during the naïve state or recovery phase following influenza infection (day 60) (Figure 6A). PCA of ~70,000 differentially methylated cytosines (DMCs, FDR *q*-value < 0.05) revealed tight clustering according to group assignment with the main variance across the dataset (PC1) reflecting methylation changes due to age (Figure 6B), consistent with prior studies (*15, 24*). We next identified genes that were both differentially expressed and had differentially methylated cytosines within their gene promoters (ANOVA, FDR *q*-value <0.05), and found 1,319 genes meeting this parameter threshold (Figure 6C). *K*-means clustering of gene expression levels revealed a substantial similarity to the DEG heat map shown in Figure 4C. Gene set enrichment analysis of these genes demonstrated that this methylation-regulated gene expression program was associated with pro-recovery processes and was significantly skewed toward young Treg cells (Figure 6C). Combined, these results show that age-related DNA methylation regulates the pro-reparative transcriptional regulatory network during recovery from influenza-induced lung injury.

**Figure 6.**
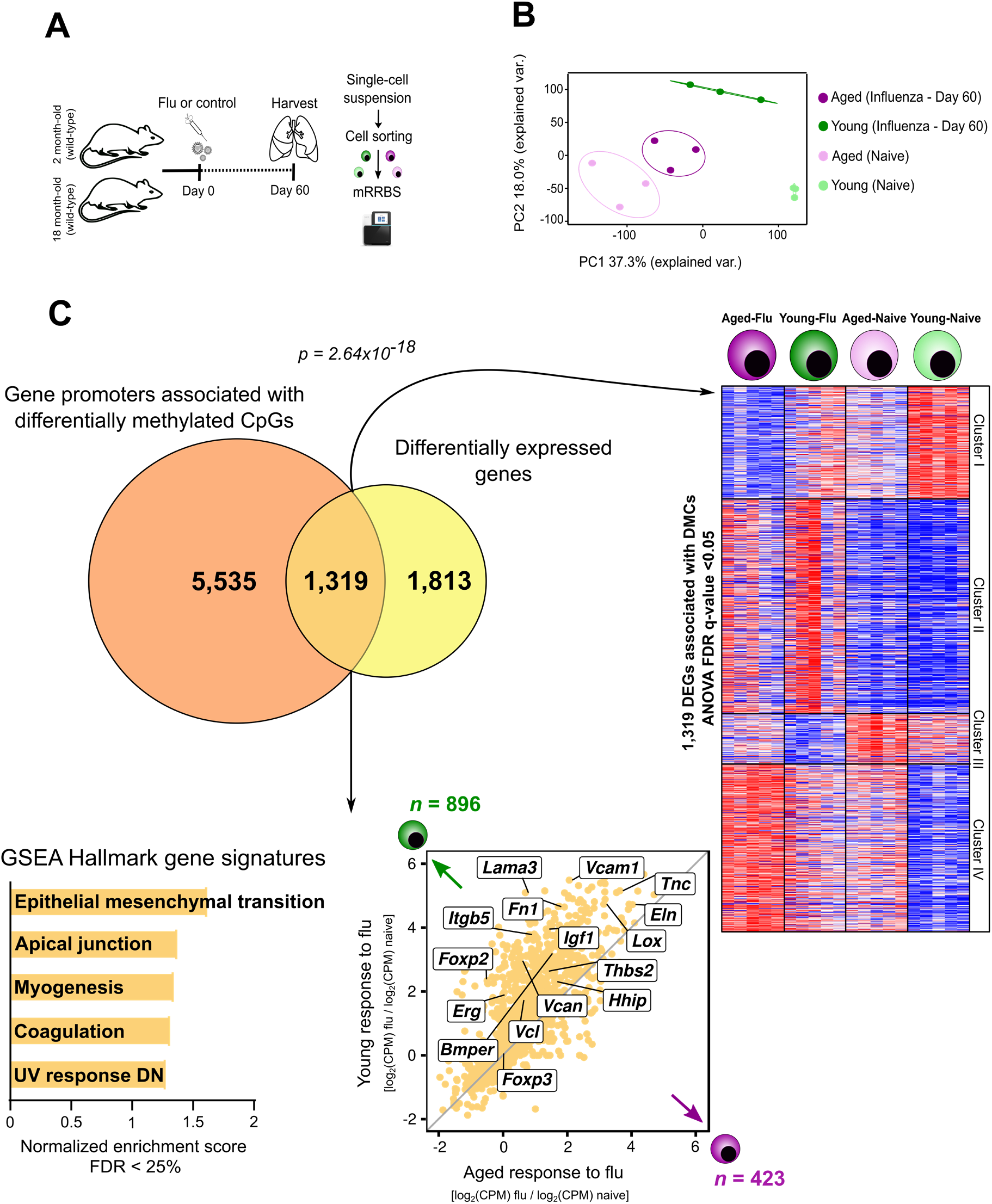
DNA methylation regulates the age-related pro-repair gene signature during recovery from influenza infection. (A) Schematic of experimental design. (B) Principal component analysis of ~70,000 differentially methylated cytosines (DMCs) identified from a generalized linear model and ANOVA-like testing with FDR *q*-value < 0.05. Ellipses represent normal contour lines with one standard deviation probability. (C) Venn diagram partitioning into differentially expressed genes or DEGs (yellow circle, 1,813 genes), gene promoters containing DMCs (dark orange circle, 5,535 genes) and genes which are both DEGs and have gene promoters containing DMCs (light orange intersection, 1,319 genes). Promoters were defined as 1 kb surrounding the transcription start site. A hypergeometric *p*-value is shown. *K*-means clustering of 1,319 genes with an FDR *q*-value < 0.05. Fold-change-fold-change plot for young Treg cell response to influenza versus aged Treg cell response to influenza infection highlighting methylation-regulated differentially expressed genes. GSEA plot with the top five positively enriched gene sets with an FDR *q*-value < 0.25. Genes were ordered by log_2_(fold-change) and ranked by the young Treg cell phenotype.

### Graphical abstract

Proposed model describing the young (**A**) and aged (**B**) pro-reparative response of Treg cells following influenza pneumonia. (**C**) Proposed model for reparative effect of heterochronic adoptive transfer of young Treg cells into an aged host.

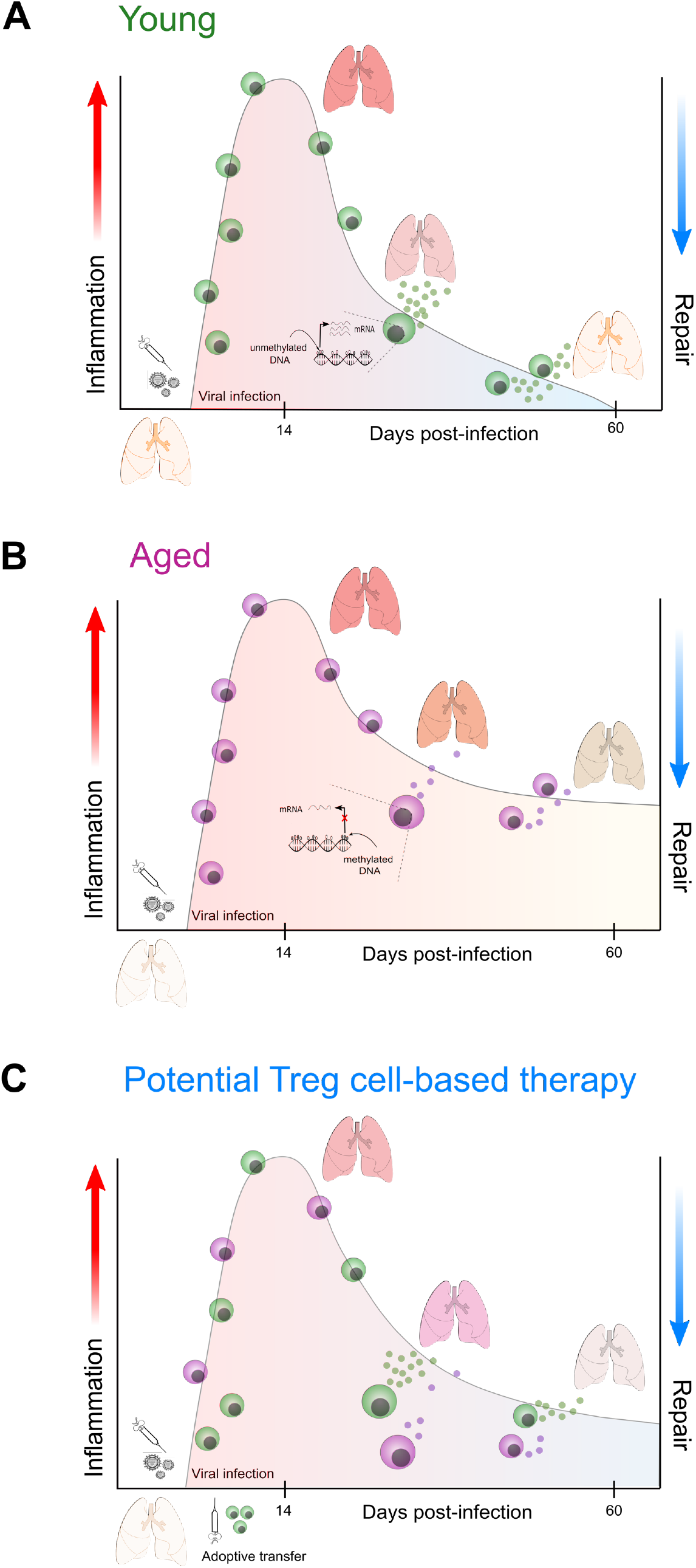

## Discussion

We sought to unambiguously address the paradigm of how aging affects Treg cell function during recovery from influenza pneumonia. We used heterochronic (age-mismatched) Treg cell adoptive transfer following influenza infection to establish that the age-related pro-repair function of these cells is determined by cell-autonomous mechanisms. While adoptive transfer of young Treg cells into aged or Treg-cell depleted hosts demonstrated a salutary effect, the transfer of aged cells into young or Treg-cell depleted hosts had a detrimental impact on mortality. Comprehensive transcriptional and DNA methylation profiling revealed age-related epigenetic repression of the youthful reparative gene expression profile and activation of maladaptive responses in lung Treg cells among aged hosts.

The ongoing coronavirus disease 19 (COVID-19) pandemic caused by the severe acute respiratory syndrome coronavirus 2 (SARS-CoV-2) represents an unprecedented challenge for the scientific community to identify novel pharmacotherapies and strategies for effective disease management. Both influenza A virus- and SARS-CoV-2-induced lung injury disproportionately affect the elderly, comprising most infection-related deaths (*1, 25, 26*). Here, we found that similar to human epidemiologic data and previous pre-clinical murine studies, aged mice exhibit increased susceptibility and impaired recovery following influenza infection. Injury to alveolar epithelial type I, II and endothelial cells disrupts the tight gas exchange barrier causing accumulation of fluid and pro-inflammatory mediators in the alveolar space, a hallmark of ARDS pathophysiology (*4*). Notably, we found that during late recovery from influenza infection, aged hosts demonstrated a decreased number of alveolar epithelial type II cells and endothelial cells when compared with young animals, suggesting that failure to repopulate the alveolar lining contributes to the observed age-related impairment in recovery. Severe influenza infection leads to a robust expansion of Krt5^+^ cells, which migrate distally to form cystic-like structures or pods intended to cover the damaged alveolar wall (*19*). These pods persist long after the initial infection, lack the capacity to generate a functional alveolar epithelium and therefore constitute an insufficient reparative response to injury (*19*). Here, we showed that aged animals display an increased percentage of Krt5^+^ cells during the recovery phase of influenza-induced lung injury, which reflects the dysregulated repair response in aged hosts.

Over the past decade, regulatory T cells have emerged as key mediators of wound healing and tissue regeneration (*6, 16, 27*). This group of specialized cells has been primarily known for their ability to suppress effector immune cell subsets leading to resolution of inflammation, but they are also capable of directly affecting tissue regeneration through production of pro-repair mediators such as amphiregulin and keratinocyte growth factor (*11, 28, 29*). Investigators have demonstrated that aging can negatively impact the composition and function of the Treg cell pool throughout the lifespan, rendering them inefficient as facilitators of tissue repair (*30*). This decline might occur through cell-autonomous mechanisms resulting in T cell maladaptive responses that lead to increased susceptibility to disease. For instance, loss of stemness accompanied by differentiation into pro-inflammatory Th1/Th17 phenotypes, activation of DNA damage responses and the senescence secretome are among some of the T cell maladaptations that result from the mounting challenges to which the T cell repertoire is exposed over a lifetime (*20*). These T cell maladaptive changes could also result from an age-related loss in stromal signals and circulating factors from the tissue microenvironment that either affect T cell function directly or render the microenvironment resistant to T cell responses. Our heterochronic adoptive Treg cell transfer experiments definitively address this paradigm, showing that the observed age-related Treg cell dysfunction is due to cell-autonomous mechanisms and dominant over the aged pulmonary microenvironment. Our data demonstrate that aging not only imparts a loss of pro-recovery Treg cell function, but also a gain of some of these maladaptive features when compared with young hosts.

What are the molecular mechanisms underpinning the age-associated Treg cell gain or loss-of pro-reparative function in the lung following influenza infection? Gene expression profiling of lung Treg cells during the recovery phase of influenza infection showed that young Treg cells significantly upregulated genes (when compared with aged Treg cells) linked to biologic processes associated with a robust pro-repair signature, including extracellular matrix organization, alveologenesis and vasculogenesis. Here, we demonstrate that the young Treg cell pro-repair program is dominated by *Areg* expression, accompanied by upregulation of IL-18 and IL-33 receptors and other genes related to the above-mentioned reparative processes. Interestingly, we found no difference when comparing the suppressive phenotype of young versus aged Treg cells, suggesting that following influenza-induced lung injury, the reparative program of Treg cells is separable and distinct from their suppressive program. This is an important observation that informs the development of novel Treg cell-based immunotherapies to specifically target molecular pathways regulating their reparative function. In regard to aged Treg cells, we found that although capable of upregulating a pro-repair program following influenza infection, it is less robust when compared with the youthful reparative response. Moreover, aged Treg cells displayed increased expression of genes associated with an effector phenotype. Accordingly, we found increased expression of Th1 canonical markers, Tbet and Ifn-γ. Whether this finding represents an age-related functional adaptability of Treg cells following influenza infection or it is the result of Treg cell lineage instability leading to effector differentiation remains unknown.

Establishment of a Treg cell specific DNA hypomethylation pattern at key genomic loci is necessary to maintain the lineage stability and immunosuppressive function of Treg cells (*13*). Epigenomic profiling has revealed that Treg cell-specific alterations in methylation patterning modulate Treg cell transcriptional programs and increase susceptibility to human autoimmune diseases (*31*). Whether epigenetic phenomena have a similar regulatory role in modulating the Treg cell reparative gene expression program remains unknown. Here, we used an unsupervised bioinformatics analysis to uncover a Treg cell-specific methylation-regulated transcriptional program enriched for reparative processes during recovery from influenza infection. Our computational integrative approach provides inferential evidence that age-related DNA methylation can modify the expression of genes linked to pro-repair processes in Treg cells but does not prove causality and therefore represents a limitation of our study. Future research could focus on leveraging epigenome editing technologies to establish the causality of age-related, Treg cell-specific DNA methylation patterns in controlling their regenerative function.

In conclusion, our study establishes that aging imparts cell-autonomous dysfunction to the reparative capacity of Treg cells following influenza pneumonia. The youthful reparative transcriptional response of Treg cells is dominated by processes linked to epithelial and endothelial cell repair and extracellular matrix remodeling, and demonstrate regulation by DNA methylation. Aged Treg cells exhibited a less robust reparative program and displayed features of maladaptive T cell responses. These findings carry important implications for the development of small molecule- and Treg cell-based therapeutics that promote restoration of lung architecture and function following viral pneumonia in our increasingly older population.

## Materials and Methods

### Mice

Young (2-4-month-old) and aged (18-22-month-old) C57BL/6 mice were obtained from NIA Aged Rodent Colony. *Foxp3^DTR^* mice were purchased from The Jackson Laboratory (Jax 016958). Animals received water *ad libitum*, were housed at a temperature range of 20-23 °C under 14/10-hour light/dark cycles, and received standard rodent chow. All animal experiments and procedures were conducted in accordance with the standards established by the United States Animal Welfare Act set forth in National Institutes of Health guidelines and were approved by the Institutional Animal Care and Use Committee (IACUC) at Northwestern University.

### Administration of influenza A virus and lung histopathology

Wild-type C57BL/6 mice were anesthetized with isoflurane and intubated using a 20-gauge angiocatheter cut to a length that placed the tip of the catheter above the carina. Mice were instilled with a mouse-adapted influenza A virus (A/WSN/33 [H1N1]) (3 pfu/mouse or 2 pfu/mouse for *Foxp3^DTR^* mice, in 50 μL of sterile PBS) as previously described (*32*).

To prepare organ tissues for histopathology, the inferior vena cava was cut and the right ventricle was perfused *in situ* with 10 mL of sterile PBS and then sutured a 20-gauge angiocatheter into the trachea via a tracheostomy. The lungs were removed *en bloc* and inflated to 15 cm H_2_O with 4% paraformaldehyde. 5-μm sections from paraffin-embedded lungs were stained with hematoxylin-eosin and examined using light microscopy with the high-throughput, automated, slide imaging system, TissueGnostics (TissueGnostics GmbH).

### Tissue preparation, flow cytometry analysis and sorting

Single-cell suspensions from harvested mouse lungs were prepared and stained for flow cytometry analysis and fluorescence-activated cell sorting as previously described using the reagents shown in Supplemental Table 1 (*33, 34*). The CD4^+^ T Cell Isolation Kit, mouse (Miltenyi) was used to enrich CD4^+^ T cells in single-cell suspensions prior to flow cytometry sorting. Cell counts of single-cell suspensions were obtained using a Cellometer with AO/PI staining (Nexcelom Bioscience) before preparation for flow cytometry. Data acquisition for analysis was performed using a BD Symphony A5 instrument with FACSDiva software (BD). Cell sorting was performed using the 4-way purity setting on BD FACSAria SORP instruments with FACSDiva software. Analysis was performed with FlowJo v10.6.1 software.

### Cytokine measurements

Lungs were harvested from young and aged mice and a single-cell suspension was obtained. Red blood cells were removed with ACK Lysis Buffer (Thermo Fisher) following the manufacturer’s instructions. Single-cell suspensions were plated on 12-well cell culture plates (Thermo Fisher) at a concentration of 1 x 10^6^ cells/mL with RPMI plus 2 μL/mL Leukocyte Activation Cocktail with GolgiPlug (BD) and incubated for 4 hours at 37 °C. After incubation, cells were resuspended in PBS and stained with a viability dye and subsequently with fluorochrome-conjugated antibodies directed at IFN-γ (clone XMG1.2), IL-17 (clone TC11-18H1) and IL-4 (clone 11B11). Data acquisition and analysis was performed as described above.

### Treg cell isolation and adoptive transfer

Splenic CD4^+^CD25^+^ Treg cells were isolated from euthanized young (2-4-month-old) and aged (18-22-month-old) C57BL/6 mice by use of magnetic separation with the EasySep™ Mouse CD4^+^CD25^+^ Regulatory T Cell Isolation Kit II (STEMCELL Technologies) according to the manufacturer’s instructions. A separate group of young and aged C57BL/6 mice were challenged with 3 pfu/mouse of influenza A virus as previously described (*32*). Twenty-four hours later a single-cell suspension of isolated 1 x 10^6^ splenic Treg cells in 100 μL of sterile PBS was obtained and transferred via retro-orbital injection into the influenza-treated mice. *Foxp3^DTR^* mice were challenged with 2 pfu/mouse of influenza A virus. Diphtheria toxin (List Biologicals, CA) was administered via intraperitoneal injection in 100μL of sterile PBS in the following doses and days relative to influenza A virus infection (day 0): 50 mcg/Kg on day −2 and 10 mcg/Kg every 48 hours starting on day 0 and ending on day 28 post-infection. Five days later, 1 x 10^6^ young or aged splenic Treg cells in 100 μL of sterile PBS were transferred via retro-orbital injection into the influenza-treated *Foxp3^DTR^* mice.

### RNA-sequencing

Flow cytometry sorted lung Treg cells were pelleted in RLT plus buffer with 2-mercaptoethanol and stored at −80 °C until RNA extraction was performed. The Qiagen AllPrep DNA/RNA Micro Kit was used for RNA and DNA simultaneous isolation (*35*). RNA quality was assessed with the 4200 TapeStation System (Agilent Technologies). mRNA was isolated from purified 1 ng total RNA using oligo-dT beads (New England Biolabs, Inc). NEBNext Ultra™ RNA kit was used for full-length cDNA synthesis and library preparation. Libraries were pooled, denatured and diluted, resulting in a 2.0 pM DNA solution. PhiX control was spiked at 1%. Libraries were sequenced on an Illumina NextSeq 500 instrument (Illumina Inc) using NextSeq 500 High Output reagent kit (1×75 cycles). For RNA-seq analysis, FASTQ reads were demultiplexed with bcl2fastq v2.17.1.14, trimmed with Trimmomatic v0.38 (to remove low quality basecalls), and aligned to the *Mus musculus* or mm10 (GRCm38) reference genome using TopHat v.2.1.0. Resultant bam files were imported into SeqMonk v1.45.4 to generate raw counts table with RNA-seq quantitation pipeline and filtered by protein-coding genes. Annotated probe reports from SeqMonk were imported into RStudio for downstream analysis. Differential gene expression analysis was performed with the edgeR v3.28.1 R/Bioconductor package using R v3.6.3 and RStudio v1.2.1578 (*36*). A genomic dataset visualization tool, Morpheus web interface (https://software.broadinstitute.org/morpheus/), was used to perform *K*-means clustering and heat maps. Gene ontology analysis was performed by using either the Molecular Signatures Database (MSigDB) from The Broad Institute or the Metascape interface. Gene Set Enrichment Analysis was performed using the Broad Institute’s GSEA v4.0.3 software with the GSEAPreranked tool. A ranked gene list from young and aged phenotypes was ordered by log_2_(fold-change) in average expression, using 1,000 permutations and the Hallmark gene set database (*37*).

### Modified reduced representation bisulfite sequencing

Genomic DNA was isolated from sorted lung Treg cells using Qiagen AllPrep DNA/RNA Micro Kit. Endonuclease digestion, fragment size selection, bisulfite conversion and library preparation were performed as previously described (*36, 38–40*). Sequencing was performed on NextSeq 500 instrument (Illumina). DNA methylation analysis and quantification were performed using Trim Galore! v0.4.3, Bismark v0.16.3, DSS v2.30.1 R/Bioconductor package and the bisulphite feature methylation pipeline from the SeqMonk platform. *Mus musculus* or mm10 (GRCm38) was used as the reference genome.

### Statistical analysis

All statistical tests are detailed under either the results section or figure legends. Statistical analysis was performed using either GraphPad Prism v8.3.0 or R v3.6.3. Computational analysis was performed using Genomics Nodes and Analytics Nodes on Quest, Northwestern University’s High-Performance Computing Cluster.

### Data and material availability

The raw and processed next-generation sequencing data sets have been uploaded to the GEO database (https://www.ncbi.nlm.nih.gov/geo/) under accession number GSE151543, which will be made public upon peer-reviewed publication.

## Acknowledgements

We thank the Northwestern University Flow Cytometry Core Facility supported by Cancer Center Support Grant (NCI CA060553). Flow cytometry Cell Sorting was performed using BD FACSAria SORP systems purchased through the support of NIH 1S10OD011996-01 and 1S10OD026815-01. Histology services were provided by the Northwestern University Mouse Histology and Phenotyping Laboratory, which is supported by P30CA060553 awarded to the Robert H. Lurie Comprehensive Cancer Center. Imaging work was performed at the Northwestern University Center for Advanced Microscopy generously supported by NCI CCSG P30CA060553 awarded to the Robert H. Lurie Comprehensive Cancer Center. We also wish to thank the Northwestern University RNA-Sequencing Center/Genomics Lab of the Pulmonary and Critical Care Medicine and Rheumatology Divisions. This research was supported in part through the computational resources and staff contributions provided by the Genomics Compute Cluster, which is jointly supported by the Feinberg School of Medicine, the Center for Genetic Medicine, and Feinberg’s Department of Biochemistry and Molecular Genetics, the Office of the Provost, the Office for Research, and Northwestern Information Technology. The Genomics Compute Cluster is part of Quest, Northwestern University’s high-performance computing facility, with the purpose to advance research in genomics.

## Author contributions

LMN and BDS contributed to the conception, hypothesis delineation and design of the study; LMN, KAH, NSM, MATA, RP, HA-V, YP and BDS performed experiments/data acquisition and analysis. LMN and BDS wrote the manuscript.

---

**Supplemental Table 1.** Flow cytometry reagents.

| **Antigen/Reagent** | **Conjugate** | **Clone** | **Company** | **Catalog number** |
| --- | --- | --- | --- | --- |
| CD25 | PE | PC61.5 | eBioscience | 12-0251-82 |
| CD25 | APC | PC61.5 | eBioscience | 17-0251-82 |
| Foxp3 | PE-Cyanine 7 | FJK-16s | eBioscience | 25-5773-82 |
| CD3ε | APC | 17A2 | eBioscience | 17-0032-82 |
| CD3ε | FITC | 145-2C11 | eBioscience | 11-0031-82 |
| CD62L | APC-eFluor 780 | MEL-14 | eBioscience | 47-0621-82 |
| CD4 | BUV395 | GK1.5 | BD | 563790 |
| CD4 | PerCP Cyanine 5.5 | GK1.5 | Biolegend | 100433 |
| CD44 | BUV373 | IM7 | BD | 612779 |
| CD44 | BV421 | IM7 | Biolegend | 103039 |
| Tigit | PerCP-eFluor 710 | GIGD7 | eBioscience | 46-9501-82 |
| CD69 | FITC | H1 2F3 | BD | 553236 |
| Helios | PE | 22F6 | eBioscience | 12-9883-42 |
| CD103 | PE-CF594 | M290 | BD | 565849 |
| Gitr | APC | DTA-1 | eBioscience | 17-5874-81 |
| Pd-1 | PerCP-eFluor 710 | J43 | eBioscience | 46-9985-82 |
| Ox40 | APC | OX-86 | eBioscience | 17-1341-82 |
| Klrg1 | BV711 | 2F1 | BD | 564014 |
| Nrp1 | PerCP-eFluor 710 | 3DS304M | eBioscience | 46-3041-82 |
| Icos | APC | C398.4A | eBioscience | 17-9949-82 |
| Tbet | PE | 4B10 | eBioscience | 12-5825-82 |
| Gata3 | PerCP-eFluor 710 | TWAJ | eBioscience | 46-9966-42 |
| Ctla-4 | APC | UC10-4F10-11 | Milipore | MABF545 |
| Ror-γ | BV421 | Q31-378 | BD | 562884 |
| IL-17 | PE | TC11-18H1 | BD | 561020 |
| IFN-γ | PE-CF594 | XMG1.2 | BD | 562303 |
| IL-4 | APC | 11B11 | BD | 562045 |
| CD45 | FITC | 30.F11 | Biolegend | 103107 |
| CD31 | PE | MEC 13.3 | BD | 561073 |
| T1α | PE-Cyanine 7 | 1.1(8.1.1) | BD | 25-5381-82 |
| CD326 | BV421 | G8.8 | Biolegend | 118225 |
| MCHII | BUV395 | 2G9 | BD | 743876 |
| Krt5 | Unconjugated<br>Alexa Fluor 488 –<br>Secondary antibody | Poly9059 | Biolegend<br>Invitrogen | 905901<br>A11039 |
| FR4 | PE-Cyanine 7 | 12A5 | eBioscience | 11-5445-82 |
| Live-dead marker | eFluor 506 | - | eBioscience | 65-0866-14 |

